# Nucleic acid sequence determinants of transcriptional pausing by human mitochondrial RNA polymerase (POLRMT)

**DOI:** 10.1101/2025.04.25.650729

**Authors:** An H. Hsieh, Tatiana V. Mishanina

## Abstract

Transcription by RNA polymerase (RNAP) lies at the heart of gene expression in all organisms. The speed with which RNAPs produce the RNA is tuned in part by the signals in the transcribed nucleic-acid sequences, which temporarily arrange RNAPs into a paused conformation unable to extend the RNA. In turn, the altered transcription kinetics determines the three-dimensional shape into which RNA ultimately folds, dictates the chromatin state, and promotes or inhibits co-transcriptional events. While pause sequence determinants have been characterized for multi-subunit RNAPs in bacteria and the eukaryotic nuclei, this information is lacking for the single-subunit RNAP of human mitochondria, POLRMT. Here, we developed a robust nucleic-acid scaffold system to reconstitute POLRMT transcription *in vitro* and identified multiple transcriptional pause sites on the human mitochondrial genomic sequence (mtDNA). Using one of the pause sequences as a representative, we performed a suite of mutational studies to pinpoint the nucleic-acid elements that enhance, weaken, or completely abolish POLRMT pausing. Finally, a search of the human mtDNA for the pause motif revealed multiple predicted pause sites, with potential roles in mitochondrial co-transcriptional processes.

## Introduction

A key component of gene expression regulation in bacteria and eukaryotes is transcriptional pausing by RNA polymerase (RNAP). These temporary, nucleic-acid sequence-specific stops by RNAPs give time for regulatory events to occur, such as interactions between transcription factors and the elongating RNAP to promote elongation, termination, or proofreading of incorrectly added nucleotides^1,2^. In eukaryotic transcription, RNAPII pausing can promote RNA processing events, such as the inclusion of alternative exons and processing of proximal poly(A) sites^3,4^. Pausing also allows co-transcriptional folding of catalytic, regulatory, or attenuator RNAs and terminator RNA hairpins^5^.

Although transcriptional pausing is a critical regulator of nuclear and prokaryotic gene expression, its mechanism and roles during transcription of the human mitochondrial DNA (mtDNA) are unknown. Mitochondria produce the bulk of cellular energy by oxidative phosphorylation (OXPHOS), with several OXPHOS protein subunits encoded in the mtDNA and transcribed by an RNAP unique to mitochondria, known as POLRMT (or mtRNAP). POLRMT initiates transcription from three known promoters within the non-coding region of mtDNA (Fig. 1A), with the help of two transcription factors, TFAM and TFB2M^6,7^. Downstream of the first light-strand promoter (LSP) are three Conserved Sequence Blocks (CSBIII, II, and I), where CSBII contains a G-rich sequence that terminates the majority of transcription events in the absence of the transcription elongation factor, TEFM^8,9^. Post-initiation and with the help of TEFM, POLRMT produces polycistronic RNA products that are co-transcriptionally processed (cleaved) into individual messenger, ribosomal and transfer RNAs (mt-mRNA, mt-rRNA, and mt-tRNA, respectively, Fig. 1A)^10^. Quantitative measurements of processed and unprocessed mtRNA levels in cells suggest that mtRNA processing is rapid and takes place largely co-transcriptionally^11^. Up to 30% of the total RNA in energy-demanding tissues is from mitochondrial transcription^12,13^, and transcription errors are linked to mitochondrial dysfunction and disease^14^. With these important processes in the mitochondria connected to transcription, the contribution of transcriptional pausing of POLRMT to their regulation is an intriguing possibility.

**Figure 1.**
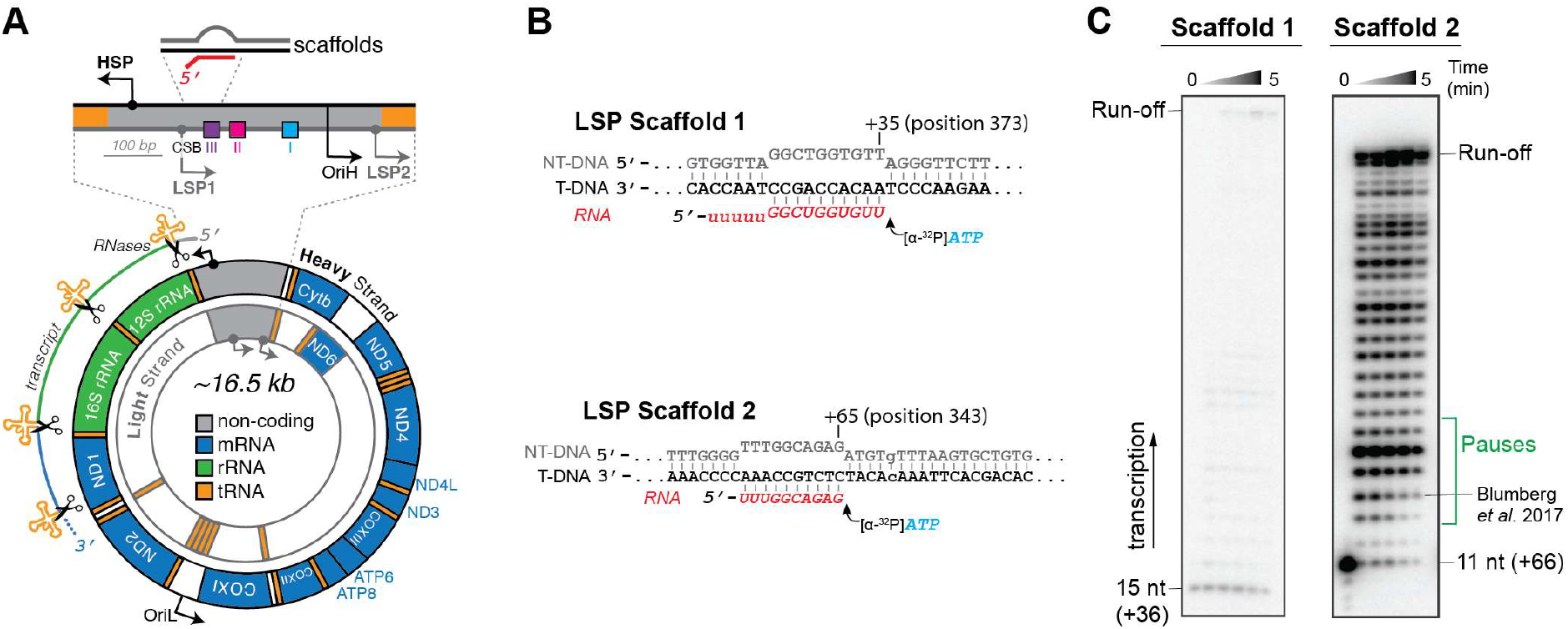
Scaffold-based transcription of mtDNA sequences. **A)** Map of the human mtDNA showing polycistronic transcription by POLRMT. Zooming in downstream of LSP1 are the CSB sites and the region of sequences we studied in scaffold-based assays. **B)** Nucleic-acid based scaffolds containing the downstream sequence of LSP1 to reconstitute POLRMT transcription *in vitro*. Scaffold 1 has more GCs at the 5’ end of the RNA and Scaffold 2 has more GCs at the 3’ end. **C)** Time course showing the difference in POLRMT transcription efficiency between Scaffolds 1 and 2.

A handful of papers concerning the pausing of POLRMT exist. In 2015, Posse et al. showed that POLRMT termination and pausing *in vitro* were reduced in the presence of TEFM, promoting POLRMT processivity and formation of long transcripts^15^. This has been followed up by Yu et al. in 2018, where pausing of POLRMT was quantified as a single-molecule level *in vitro* by magnetic tweezer experiments, and TEFM was shown to decrease the duration and frequency of long-lived pauses likely involving the formation of nascent RNA secondary structure^16^. These results are consistent with other literature showing TEFM’s role in promoting processivity *in vivo*^17,18^. Finally, *in vivo* nascent RNA sequencing by global run-on (GRO-seq) and precision run-on sequencing (PRO-seq) in 2017 suggested pausing occurring near CSBIII on the mtDNA^19^, although at a low resolution. The mechanism and sequence determinants of POLRMT pauses, however, remain unknown.

In this work, we established a robust protocol for reproducible reconstitution of POLRMT transcription *in vitro* by employing scaffold-based assays with the SC1 RNA sequence used in RNA structure probing field^20–22^. Using this system, we captured transcriptional pausing of POLRMT with single-nucleotide resolution on previously difficult-to-transcribe sequences, such as the consensus elemental pause and sequences downstream of the mitochondrial light strand promoter (LSP). Using a representative pause scaffold, we then characterized the nucleic acid sequence determinants of POLRMT pausing by systematic mutational studies.

## Materials and Methods

### Proteins

The plasmids for POLRMT (pPROEXHTb-6xHis-POLRMT (amino acids 43-1230)) and TFAM (pPROEXHTb-6xHis-TEV-TFAM (amino acids 43–246)) were gifts from Prof. Smita Patel (Rutgers University). The TFB2M plasmid (pT7TEV-HMBP4-TFB2M (amino acids 21-396)) was a gift from Prof. Miguel Garcia-Diaz (Stony Brook University School of Medicine). A detailed purification protocol for POLRMT, TFAM, and TFB2M can be found in our Bio-protocol paper^7,23^. *E. coli* RNAP purification details were also described previously^24^.

Expression construct for TEFM (residues 36-360, with C-terminal 6xHis tag in pET21 vector) was a gift from Prof. Dmitry Temiakov (Thomas Jefferson University)^25,26^. We used the protocol from Hillen et al. as a starting point^26^. Specifically, *E. coli* BLR (DE3, recA^-^) cells transformed with TEFM-encoding plasmid were grown in 2xYT media overnight at 30 °C. 5 mL of this starter overnight culture was inoculated into each liter of 2xYT and shaken at 200 rpm at 37 °C for 3-4 hrs. When OD_600_ reached 0.4 units, the temperature was lowered to 16 °C. 0.2 mM of IPTG (final concentration) was added to induce TEFM expression, and the cultures were shaken overnight at 16 °C. Cell pellets were harvested and lysed by sonication. TEFM was purified from the crude lysate through Ni Sepharose pre-packed column (HisTrap HP 5 mL, GE Healthcare) in 5-500 mM gradient of imidazole (over XX column volumes), and then Heparin Sepharose pre-packed column (HiTrap Heparin HP 5 mL, GE Healthcare) in 250-1500 mM gradient of NaCl (over XX column volumes). The elution fractions were further passed through the size exclusion column (Superose 6 Increase 10/300 GL). Pure TEFM was concentrated to 32 μM with a 3K MWCO Amicon concentrator, aliquoted and stored at −80 °C in Storage Buffer (10 mM Tris-Cl, pH 7.9, 25% glycerol, 200 mM NaCl, 100 μM EDTA, 1 mM MgCl_2_, 10 mM DTT).

### Nucleic Acids

All oligonucleotides used are listed in Table S1. DNA and RNA oligos were obtained from Integrated DNA Technologies. All single-stranded RNA and DNA used in experiments with incorporation labeling by [α-^32^P]ATP (Revvity) were purified by denaturing PAGE, gel extraction, and ethanol precipitation (detailed protocol described previously^24^). We have observed that POLRMT will label the linear double-stranded DNA template when incorporation labeling by [α- ^32^P]ATP is performed, resulting in bands greater than the expected RNA run-off size and consistent with the length of the DNA template instead. The DNA used in experiments with [γ-^32^P]ATP (mutational studies) did not need to be purified, as only the RNA bands will appear on the gel.

In this paper, we use “LSP” to refer to LSP1 of the mtDNA and its downstream sequence. As a side note, when we use a template DNA containing the G-rich CSBII region, we are unable to resolve bands around that region and see samples stuck near the top of our gels, even under various denaturing conditions. That is why we decided to use a template sequence that ends before the CSBII region of LSP.

### Scaffolds-Based Transcription with Incorporation Labeling of RNA

The SC1 RNA-based scaffolds used in the experiments in Figs. 2-4 (Scaffolds 3 and 4 in the SI), were reconstituted by the following methods, as previously described with modifications^24,27,28^. First, 10 μM of the template DNA (T-DNA) was hybridized to 5 μM SC1-RNA in UltraPure DNase/RNase-Free Water (Invitrogen) in a thermocycler by heating at 95 °C for 3 min, 75 °C for 2 min, rapidly cooling to 45 °C, and then cooling to room temperature by 1 °C/min. 0.5 μL of RNase Inhibitor Murine (New England Biolabs) was added to the scaffold (T-DNA/RNA hybrid) and was stored at −20 °C until use. Before the transcription experiment, the 10x EC was reconstituted by mixing 1.5 μM of RNAP (either POLRMT or *E. coli* RNAP) and 0.5 μM of the scaffold in the following Transcription Buffer based off of Ramachandran et al^28^: 50 mM Tris-acetate (pH 7.9), 70 mM potassium glutamate, 5 mM Mg acetate, 0.05 % Tween-20, 1 mM DTT (see note below about potassium glutamate in buffer). After incubating at 37 °C for 10 min, 1.5 μM non-template DNA (NT-DNA) was added to complete the EC assembly and incubated at room temperature for another 10 min. The ratio of the components was RNA : T-DNA : NT-DNA : RNAP = 1 : 2 : 3 : 3.

**Figure 2.**
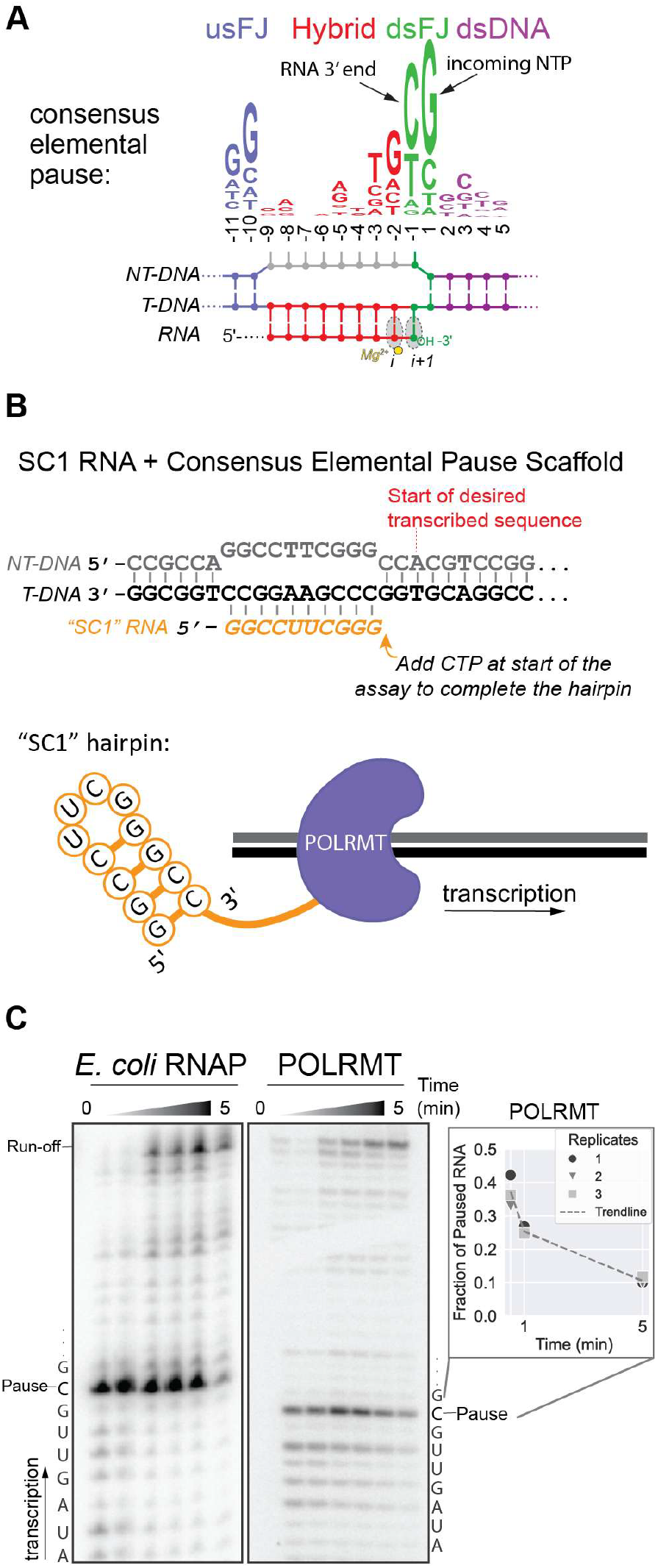
Transcription through the consensus elemental pause sequence. **A)** The consensus elemental pause sequence conserved in various organisms. **B)** Design of the SC1 RNA-based transcription on a scaffold. **C)** Time course showing both *E. coli* RNAP and POLRMT recognize the consensus elemental pause sequence.

**Figure 3.**
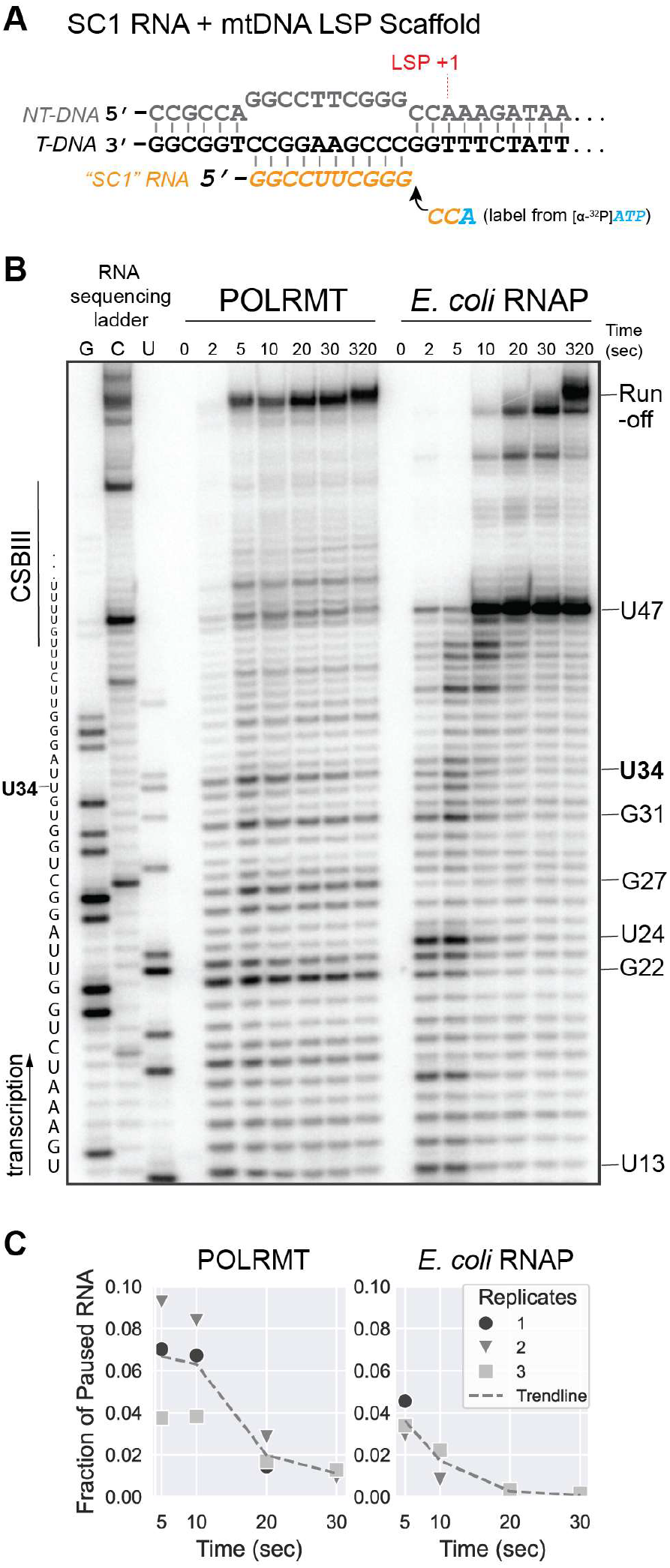
Transcription through mtDNA sequence. **A)** Scaffold containing the SC1-RNA sequence and region downstream of LSP1 of the mtDNA. **B)** Time course showing both *E. coli* RNAP and POLRMT transcribe and pause on the region downstream of LSP1. **C)** Quantification of the pause fraction at U34.

**Figure 4.**
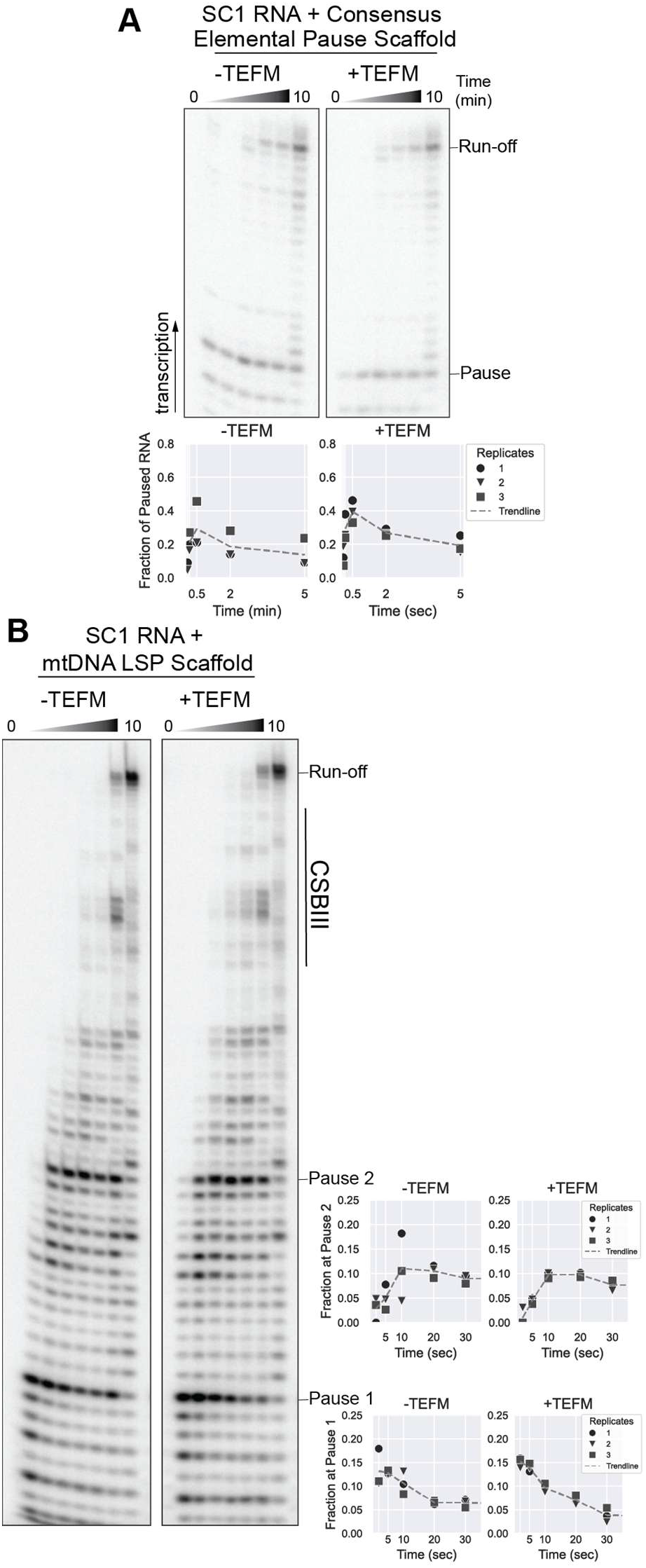
Effect of TEFM on POLRMT pausing. **A)** Time course of POLRMT transcribing the consensus elemental pause sequence ±TEFM. **B)** Time course with the region downstream of LSP ±TEFM, with quantification of two representative pauses.

Next, the ECs were diluted from 10x to 4x in the Transcription Buffer, and 0.1 mg/ml of heparin (Fisher) was added to bind any free RNAP in solution. The ECs were incubated for 3 min at room temperature before starting transcription elongation reactions. Then, 200 μM (4x of the final NTP concentration) of CTP (High Purity rNTPs (Cytiva)), trace amount of [α-^32^P]ATP (Revvity), and Transcription Buffer were added to the ECs to dilute them from 4x to 2x. Reactions were incubated for 1 min at room temperature to transcribe the full SC1 RNA hairpin and radiolabel the RNA. One sample (5 μL) was taken out to mix with an equal volume of Transcription Buffer, and then quenched with an equal volume of 2x Stop Buffer (90 mM Tris– borate buffer, 8 M urea, 47.5 mM EDTA, 50% formamide, and 0.02% of both xylene cyanol and bromophenol blue) for the zero timepoint. Similar steps were repeated for the timepoints (2 s, 5 s, 10 s, 20 s, 30 s, and 5 min) where 5 μL EC samples were taken out to mix with an equal volume of 100 μM of ATP, UTP, and GTP (for final NTP concentration of 50 μM), and then the reactions were stopped at those times with an equal volume of 2x Stop Buffer. In the reactions with TEFM, an additional “chase” timepoint was collected where 5 μL EC sample was mixed with an equal volume of 600 μM ATP, UTP, GTP, and CTP (for final concentration of 300 μM), and the reactions were stopped after 10 min with an equal volume of 2x Stop Buffer. The chase reactions were also performed to make the RNA sequencing ladder. Individual 3’-deoxynucleotides (1000 μM) (TriLink BioTechnologies) were used in a 10:1 ratio with all four rNTPs (100 μM) in 10 min chase reactions with *E. coli* RNAP on the SC1 RNA scaffold containing the downstream sequence of LSP.

To resolve the RNA species by a denaturing PAGE, the samples were heated for 2 min at 95 °C and loaded onto a 10% polyacrylamide (19:1 acrylamide:bisacrylamide), 8 M urea gel in TBE (Fisher). The gel was dried and exposed overnight to a PhosphorImager screen, which was imaged by a Typhoon PhosphorImager.

The scaffolds used in the experiment in Figs. 1 (Scaffolds 1 and 2 in the SI) were reconstituted by methods similar to described above with the following differences: Buffer follows recipe from Ramachandran et al.^28^: 50 mM Tris-acetate (pH 7.9), 50 mM potassium glutamate, 10 mM Mg acetate, 0.05 % Tween-20, 1 mM DTT. No RNase Inhibitor Murine was added. 10x EC concentrations were incubated at 37 °C for 15 min, and after the NT-DNA was added, ECs were incubated at 37 °C for an additional 10 min. The ECs were incubated with just [α-^32^P]ATP for 1 min. Then, timepoints were collected at 0, 2 s, 6 s, 10 s, 1 min with 100 μM of all four rNTPs (for final concentration of 50 μM) and 5 min chase with 500 μM of all four rNTPs (for a final concentration of 250 μM).

In addition, we have observed that POLRMT pausing is sensitive to salt concentration in our buffer, where we use potassium glutamate. For our scaffold assays, pauses and overall band intensity are optimal at 100 mM total salt in the reaction, including the storage buffer of the proteins. Potassium glutamate is used in pausing experiments to reduce the effects of chloride ions on pause escape while also resembling cellular environment more closely^29–32^. Potassium glutamate is prone to degradation in solution over time^33^, so it is best to use freshly made transcription buffer.

### Scaffolds-Based Transcription with 5’-end Labeling of RNA

The scaffolds used in the experiments in Figs. 5, 6, and S2 (Scaffolds 5-14 in the SI) were reconstituted by methods similar to described above for the SC1 RNA-based scaffolds, but the RNAs were first 5’-end labeled with ^32^P. Specifically, 20 μM PAGE-purified RNA, T4 Polynucleotide Kinase (PNK; New England Biolabs), [γ-^32^P]ATP (Revvity), and 1x PNK buffer were incubated at 37 °C for 30 min. The reaction was heated for 3 min at 95 °C to inactivate PNK, and the labeled RNA was annealed to the T-DNA, as described above. After incubation with POLRMT and NT-DNA, the unbound labeled RNA was removed from the 10x ECs with an Illustra Microspin G50 column (Cytiva). The ECs were diluted to 2x with the Transcription Buffer (50 mM Tris-acetate, pH 7.9, 70 mM potassium glutamate, 5 mM Mg acetate, 0.05 % Tween-20, 1 mM DTT) and incubated with heparin. Reactions were stopped at varying timepoints and no “chase” timepoint was collected. Samples were run on a 15% polyacrylamide (19:1 acrylamide:bisacrylamide), 8 M urea gel in TBE. The gel was dried, exposed overnight, and imaged as described above

**Figure 5.**
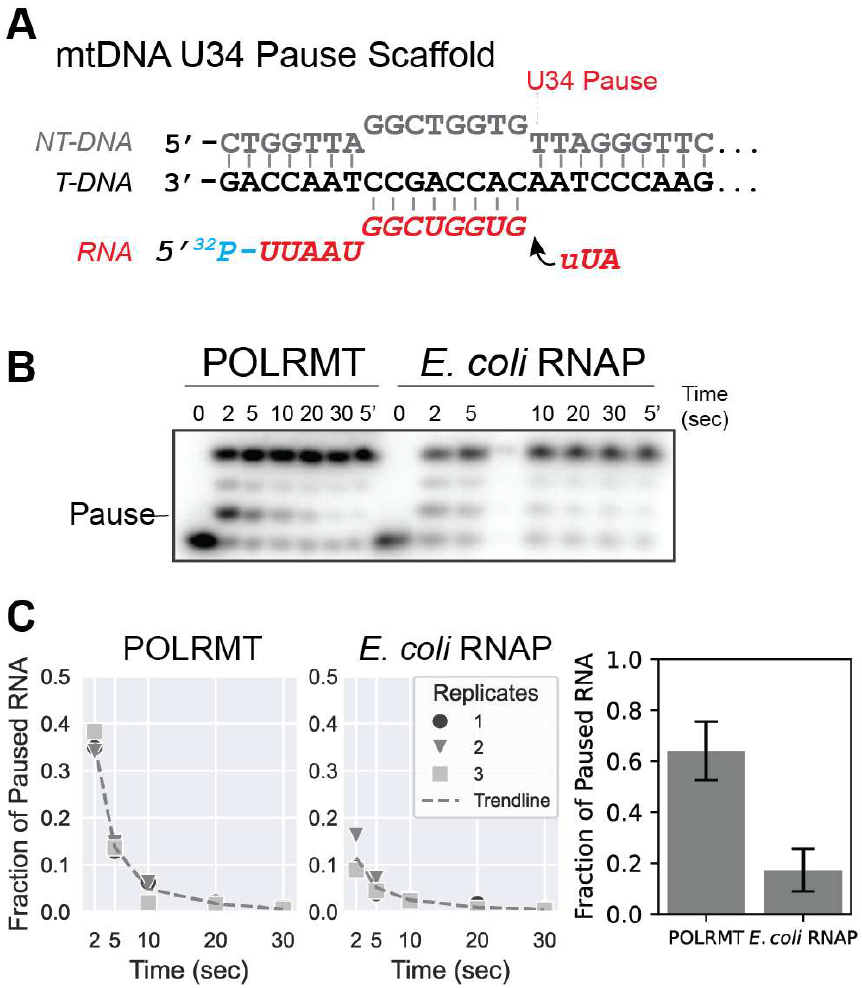
Representative POLRMT elemental pause. **A)** Scaffold starting 1 nt before the pause site observed at U34 downstream of LSP. **B)** Time course of POLRMT and *E*. coli RNAP transcribing the U34 pause scaffold. **C)** Quantification of the fraction at the pause, showing that POLRMT intrinsically pauses on this sequence.

**Figure 6.**
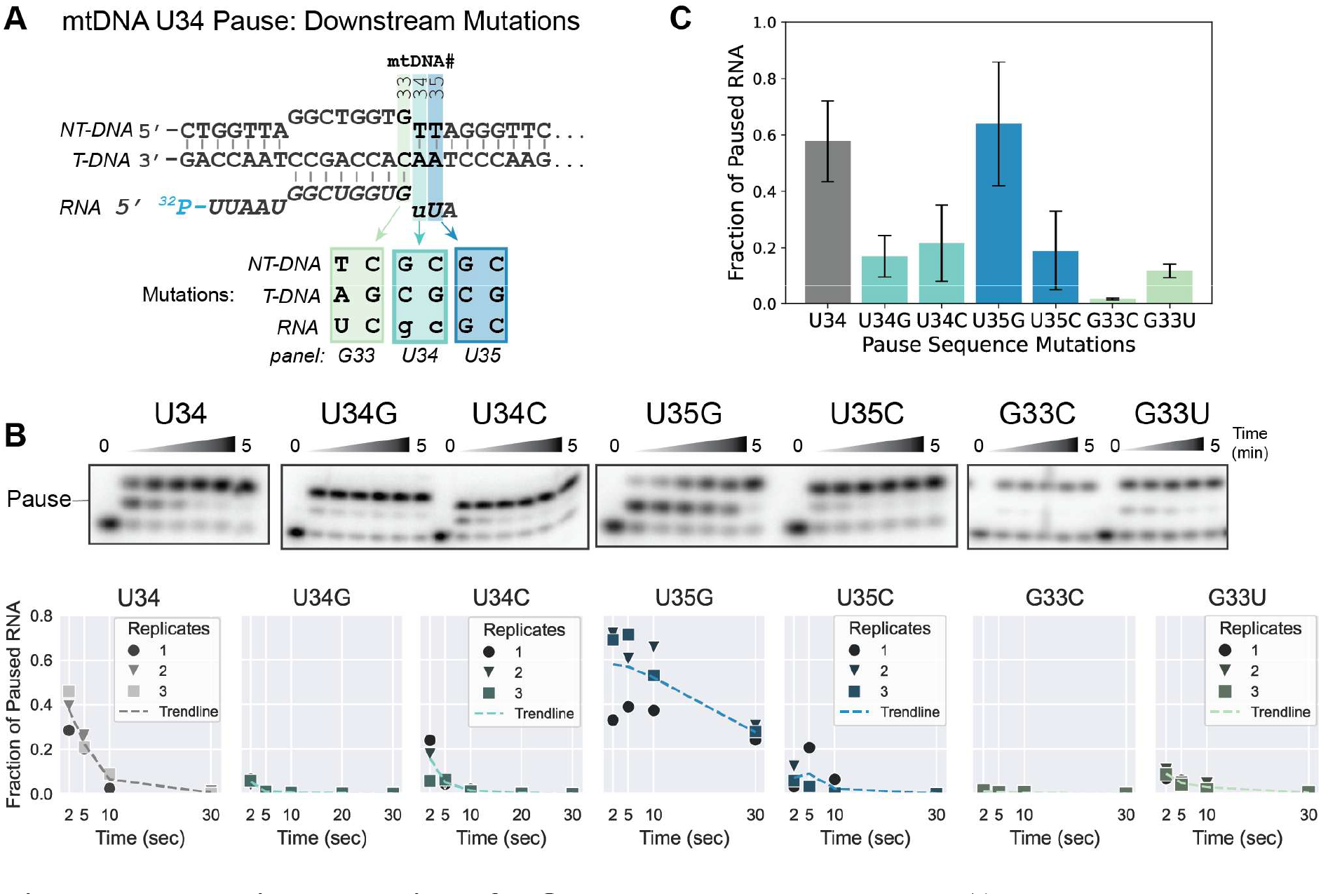
Mutational studies of POLRMT elemental pause. **A)** U34 pause sequence showing the mutations to the sequence at the downstream fork: the pause site (U34) and the nucleotides before (G33) and after it (U35). **B)** Time course of the U34 (WT) pause scaffold and each mutation, with quantification of the pause fraction over time. **C)** Bar graph showing the fraction of POLRMT/RNA at the pause for each pause sequence mutation. Error bars are are the standard deviation of the pause fraction from data collected in triplicate.

### Data Quantification

The fraction of RNA at the pause was quantified for each time point using ImageQuant TL analysis software (Cytiva). The plots of pause fraction over time were created in Python 3 using Matplotlib and Seaborn. For the mutational studies (Fig. 6), an exponential decay function was fit to the data with Python^34^. The pause fraction of each plot was calculated by finding the y-intercept and plotted as a bar graph (Fig. 6C). The pause RNA fraction graphs report the average and standard deviation of time points performed in triplicate.

## Results

### Reproducible reconstitution of POLRMT pausing *in vitro*

To visualize transcription kinetics with single-nucleotide resolution, we used nucleic-acid scaffold-based assays, where POLRMT starts transcription on a short, pre-synthesized strand of RNA (Fig. 1B). This allows POLRMT to bypass the initiation phase to elongation, where transcription is more robust. We screened scaffolds containing the sequence after the LSP, ending between CSBIII and CSBII – close to the potential pauses observed in the *in vivo* sequencing by Blumberg *et al*^19^. From screening these scaffolds, we observed strikingly different transcription efficiencies depending on the G:C pair distribution within the starting RNA:DNA hybrid. Specifically, G:C base pairing at the 3’ end of the hybrid supported robust transcription, as seen in Scaffold 2 (Fig. 1C). We concluded that having more G:C pairs at the 3’ end stabilizes the RNA:DNA hybrid, promoting productive POLRMT binding. This is consistent with prior scaffold-based transcription work showing POLRMT robustly extending a GC-only RNA^35^.

The high sensitivity of POLRMT to the GC content and particular distribution of G:C base pairs within the RNA:DNA hybrid makes choosing the optimal sequence from the mtDNA for a scaffold time-consuming. To have POLRMT transcribe every time on a scaffold sequence, we drew inspiration from the “SC1” RNA sequence, which is commonly used in the RNA structure probing field and contains the sequence features favorable for productive POLRMT reconstitution^20,21^. The idea is that POLRMT will efficiently bind to the GC-rich SC1 RNA-based scaffold and after transcribing the full SC1 RNA sequence, the RNA will fold itself into a short hairpin (Fig. 2B). This SC1 hairpin has experimentally been shown not to affect the downstream RNA in structure probing experiments^22^, which is important because nascent RNA folding is known to modulate pausing in other RNAPs^36,37^.

POLRMT will then continue to transcribe customized sequences of interest inserted after the universal starting SC1 RNA, without the need to optimize and troubleshoot the scaffold features every time. The following two sections demonstrate the utility of this SC1 RNA-based scaffold transcription approach for POLRMT.

### POLRMT recognizes the consensus elemental pause sequence identified in bacteria and the nucleus

We first used the SC1 RNA-based scaffold to see if POLRMT pauses on the *E. coli* RNAP consensus elemental pause sequence (Fig. 2A). This consensus sequence contains the most common nucleotides (nt) on which *E. coli* RNAP pauses as identified by native elongating transcript sequencing (NET-seq) *in vivo*^38,39^. These sequence elements are recognized as a pause signal by various bacterial RNAPs and mammalian RNAPII. *E. coli* RNAP pausing on the consensus elemental pause has been extensively characterized *in vitro* ^40,41^. While the molecular mechanisms of “sensing” the nucleic-acid sequence motif by RNAP are still being investigated, it is known that transcribing the consensus elemental pause sequence induces a conformational change in *E. coli* RNAP (“swiveling”) that blocks the addition of the incoming nucleotide to the growing RNA strand ^41^. The extent and propensity of bacterial RNAP to swivel, and thus pause, are modulated by transcription factors and folded nascent RNA^36,42–46^.

Initially, we used the same consensus elemental pause scaffold sequence as in the *E. coli* RNAP studies but did not see any RNA extension with POLRMT. Similar to the issues with our Scaffold 1 described above, it was likely due to the lack of G:C pairs in the 3’-proximal half of the RNA:DNA hybrid. In stark contrast, with the SC1 RNA sequence as the starting RNA and the consensus elemental pause sequence downstream of it, both *E. coli* RNAP and POLRMT were able to transcribe and pause at the expected site (Fig. 2C). POLRMT thus recognizes the *E. coli* consensus elemental pause sequence, indicating that the mitochondrial RNAP pausing mechanism may resemble that of bacterial and eukaryotic RNAPs.

### POLRMT pauses on the mtDNA sequence downstream of LSP

Next, we investigated POLRMT pausing on the human mtDNA sequence. We previously had issues with promoter-based assays resulting in low signal intensity (Fig. S1), so we applied the SC1 RNA-based scaffold approach to study POLRMT transcription through the downstream region of the LSP (Figs. 1A and 3A). Specifically, we focused on the mtDNA sequence up to CSBII, which is relatively AU-rich.

We saw robust transcription by POLRMT and observed multiple pausing sites in the LSP-proximal stretch of mtDNA (Fig. 3B). *E. coli* RNAP transcription reactions were included as well in the experiment as an example of a well-characterized RNAP that pauses. Both POLRMT and *E. coli* RNAPs pause, but not all at the same locations. *E. coli* RNAP appears to pause more frequently but for shorter amounts of time, whereas POLRMT seems to dwell on fewer pause sites but for longer. We picked a representative pause site (U34) to quantify, observing a higher fraction of POLRMT at the pause compared to *E. coli* RNAP (Fig. 3C).

Aside from temporary pauses, we also observed multiple termination or permanent arrest sites for both RNAPs. For instance, *E. coli* RNAP appeared to terminate at U47 (361 of the mtDNA), where it transcribed an 8 nt U-rich region followed by three Gs. U-tracts are known to cause backtracking or intrinsic termination of *E. coli* RNAP^1,47^, which may be causing the termination bands. POLRMT on U47, on the other hand, appears to have a small fraction pausing at the site rather than a large fraction terminating or backtracking like *E. coli* RNAP.

### TEFM does not appear to affect intrinsic pausing of POLRMT

We repeated the consensus elemental pause and the LSP experiments above with TEFM. On the consensus elemental scaffold, no significant changes were seen in the presence of TEFM, where POLRMT was expected to pause intrinsically (Fig. 4A). On our LSP scaffold with the expected run-off of ~84 nt (ending at +72 of LSP, before CSBII), we also did not see significant differences with POLRMT entering or escaping the pauses ±TEFM (Fig. 4B). This is consistent with prior *in vitro* work showing that the longer the template (100-3000 nt, especially with the CSBII region included), the greater the effects of TEFM on POLRMT ability to produce the full-length transcripts^15,17^. In addition, Yu et al. observed that TEFM decreased the duration and frequency of long-lived pauses (>4 sec) likely involving RNA secondary structure formation in the nascent transcript, but not the short-lived pauses^16^. It is thus possible that the lack of RNA structure formation in our pause experiments explains why we did not see much change in the presence of TEFM, indicating that TEFM does not regulate intrinsic (RNA-structure independent) pausing of POLRMT.

### Sequence determinants of POLRMT pausing

We chose the POLRMT pause site U34 downstream of LSP (position 375 on the human mtDNA) as a representative pause sequence (as described above in Fig. 3C). We chose this site because (i) we consistently observed pausing at this site in multiple biological replicates, and (ii) POLRMT efficiently extends a scaffold containing the U34 pre-synthesized RNA sequence, without the need for the SC1 RNA. Here, we reconstituted POLRMT elongation complex (EC) 1 nt before the pause, allowing precise and quantitative measurement of POLRMT flux through the pause site. Furthermore, because the starting RNA is short (13 nt) and cannot form secondary structures within the paused EC (PEC), any pausing observed must be due to the intrinsic propensity of POLRMT to recognize the pause sequence, i.e., elemental pausing. We saw that POLRMT pausing at the U34 site was preserved when starting 1 nt before the pause (Fig. 5B), with about 64% of POLRMT molecules pausing (Fig. 5C). This indicates that POLRMT intrinsically pauses on this sequence, not as a result of nascent RNA folding.

To determine whether POLRMT elemental pausing is sequence specific like bacterial RNAP and RNAPII, we systematically mutated the U34 pause scaffold in the regions shown to be critical to multi-subunit RNAP pausing and assessed POLRMT pausing behavior (Fig. 6A). First, we tested if the nature of the nucleobase (pyrimidine vs. purine) at the pause site U34 (−1 position) matters (Fig. 6B, left panels). The U-to-G and U-to-C substitutions greatly reduced pausing from 58% to about 20% (Fig. 6C). This suggests that the pyrimidine at the 3’-end of RNA transcript is important for POLRMT to pause, which is also the case for the consensus elemental pause in bacterial and nuclear genomes.

Second, we mutated U35, the incoming NTP site (+1 position, Fig. 6B, middle panels). Surprisingly, the U-to-G mutation increased the pause fraction, from 58% to 64%, and the dwell time, making this the “strongest” pause sequence in our experiments. This is consistent with the consensus elemental pause sequence, where G at the +1 position is the dominant nucleobase observed with *in vivo* paused multi-subunit RNAPs^39^. The POLRMT pausing fraction decreased to about 20% on the U-to-C scaffold, similar to the U34 mutations.

We next mutated G33, the nucleotide before the pause (−2 position, Fig. 6B, right panels). G mutated to C almost completely abolished pausing, decreasing the pause fraction to less than 2%. Transcription of the G-to-U scaffold showed the second most reduction to 13%. This shows that G at the −2 position is needed for strong pausing of POLRMT, which is again consistent with the consensus pause elements, where a G at the −2 position within the RNA:DNA hybrid is expected to produce a stronger pause.

Finally, we looked at the upstream fork junction of the PEC, where G at the −10 position appears to be an important nucleobase for RNAP pausing *in vivo* and *in vitro* (Fig. S2). The U34 mtDNA PEC has an A at this position. While we observed that the A-to-G substitution almost halved the pause fraction, it was still a substantial amount of pausing at 44% (Fig. S2B,C). This suggests that a purine at −10, especially an A, is sufficient to support POLRMT pausing.

By aligning all the nucleic-acid pause sequences we observed along with these substitutions, we noted the following trends: 1) POLRMT is able to pause on U, G, or C, but the pause fraction appeared to be highest with U at the pause site; 2) U and G are strongly preferred at the incoming nucleotide site; 3) G at the −2 position seems essential for a strong pause, and 4) A is preferred at the −10 position but G supports pausing as well. With these results, we constructed a consensus pause motif expected to cause strong pausing for POLRMT: 5’_-10_ RNNNNNNNGTG_+1_ 3’, where R is A or G, and N is any base.

## Discussion

In this work, we studied the uncharacterized pausing behavior of POLRMT with scaffold-based ensemble *in vitro* transcription assays. We first designed a universal starting RNA:DNA duplex encoding an SC1 hairpin, which thanks to its GC content and GC distribution, favored reproducible POLRMT binding to the scaffold and RNA extension (Fig. 2B). Appending a customized DNA sequence downstream of this starting duplex allowed reproducible POLRMT transcription for end-point and time-resolved measurements of RNA elongation. Furthermore, this design enabled us to study POLRMT transcription kinetics on previously hard-to-transcribe sequences (e.g., AU-rich). Having such a working *in vitro* system can help the field gain insight into questions about pausing occurring inside cells, such as location and duration^48^.

Using the above universal scaffold design, we showed that POLRMT recognizes the *E. coli* consensus elemental pause sequence (Fig. 2C). This sequence is conserved in other prokaryotic and eukaryotic genomes and may indicate that POLRMT, a single-subunit phage-like RNAP, has a similar mechanism of transcriptional pausing to multi-subunit RNAPs. The details of this mechanism, such as the global and active-site conformation of POLRMT and nucleic acids, will be the subject of future work. We also used the SC1 RNA scaffold design to monitor POLRMT transcription through the region downstream of the human mtDNA LSP and observed multiple pauses by POLRMT (Fig. 3B).

In *E. coli*, the transcription elongation protein NusG serves as an anti-pausing factor by preventing *E. coli* RNAP swiveling^49–51^. This to us raised the question of whether TEFM, a transcription elongation factor in human mitochondria, would play a similar role in precluding POLRMT from arranging into a paused conformation. When we performed the consensus elemental pause experiments with TEFM, however, we observed little change in POLRMT paused fraction, pause duration, or location (Fig. 4A). Similar results were seen when the experiment was repeated with the template containing the downstream region of LSP of the mtDNA, indicating that TEFM may not inhibit intrinsic (or elemental) pausing of POLRMT (Fig 4B). The mechanism of POLRMT intrinsic pausing remains uncharacterized, and it is unknown if POLRMT pauses are modulated by nascent RNA folding like in multi-subunit RNAPs. Future experiments may determine if POLRMT rearranges into different types of paused conformations, such as those in the presence of folded nascent RNA, and if the kinetics of entry into and escape from those pauses are affected by TEFM. In addition, TEFM has been shown to help POLRMT transcribe long stretches of DNA (about 3000 nt by Posse *et al*. (2015) *in vitro*), particularly through the G-rich, CSBII region which could produce structured RNA – an observation we reproduced (Fig. S1.C)^15,25,52^. Thus, it is possible that TEFM could play a role in structured RNA-mediated POLRMT pauses.

Although both *E. coli* RNAP and POLRMT recognized the consensus elemental pause (Fig. 2C), a large fraction of *E. coli* RNAPs paused at the consensus site (~90%) but a much smaller fraction of POLRMT molecules (~40%) paused at this site. Likewise, on the sequence downstream of mtDNA LSP, both RNAPs paused but not at the same sites or for the same amount of time (Fig. 3B), suggesting RNAP-specific attributes of pausing on nucleic-acid sequences. We thus next attempted to determine if POLRMT has its own “consensus elemental pause” sequence, by systematically mutating the sequence elements of a representative POLRMT pause (LSP +34 or position 375 in the reference human mtDNA genome, Fig. 5).

We first confirmed that POLRMT still paused at the U34 site without the upstream nascent RNA, by positioning POLRMT on a short pre-synthesized RNA with its 3’ end one nucleotide before the expected pause site (Fig. 5A). We indeed saw POLRMT pause at U34, indicative of an “elemental” pause at this site that is not caused by nascent RNA folding (Fig. 5B). We then performed mutational studies on this representative POLRMT pause sequence, by introducing single-nucleotide substitutions at key positions of the transcription complex, specifically the 3’ active-site nucleotides and 5’ RNA:DNA hybrid base pairs (Figs. 6 and S2). Briefly, we found that 1) pyrimidine at the pause site (position +34 of LSP) is preferred for pausing, with U having higher pause fraction over a C; 2) a G at the +1 site (position +35) – the incoming NTP to be added upon pause escape – significantly prolonged the pause; 3) a G at the −2 position (position +33) appeared essential for pausing, and 4) purines at the −10 position upstream, with A preferred over G, supported POLRMT pausing.

Combining the mutational data, we created a consensus pause motif that we predict to produce a long-lived POLRMT pause on mtDNA (Fig. 7A). We searched both strands of human mtDNA for the consensus pause motif (5’ _-10_RNNNNNNNGTG_+1_ 3’, where R is A or G, and N is any base). We found 23 matches on the strand encoding the heavy-strand genes (Fig. 7B) and over 200 matches for the strand encoding the light-strand genes. Note that this searches for only the “strongest” pause sequence we observed in our assays; other combinations of nucleic acid sequences may produce weaker pauses that this search would miss. The pauses we found on the heavy strand are scattered, with two in the non-coding region and others within rRNAs, mRNAs, and tRNAs (Fig. 7B). We hypothesize that these pauses may aid in co-transcriptional RNA folding and/or other co-transcriptional events, analogously to the characterized roles of RNAP pausing in bacteria and the eukaryotic nuclei.

**Figure 7.**
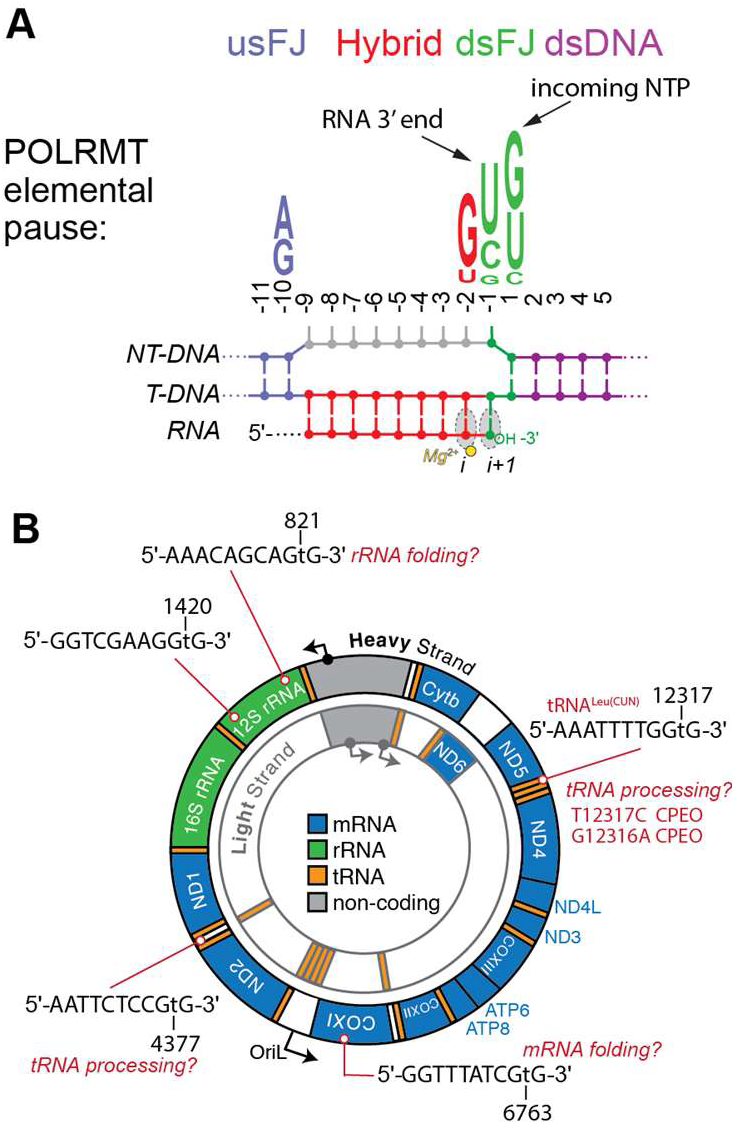
Examples of predicted POLRMT elemental pause sites on mtDNA. **A)** “Consensus elemental pause” for POLRMT determined by the mutation studies. **B)** Search of the mtDNA for this pause motif yielded potential pause sites that could play roles in regulating co-transcriptional RNA processing or nascent RNA folding.

POLRMT pausing altered by mtDNA mutations could have adverse effects on RNA folding into biologically functional shapes or RNA processing and may be linked to disease. For example, a pathogenic mutation at T12317C, a potential pause site on tRNA^Leu(CUN)^, is linked to symptoms including chronic progressive external ophthalmoplegia (CPEO), myopathy, and diabetes^53,54^. This predicted pause is in the T-loop of the tRNA, where POLRMT pausing could allow folding of the tRNA or permit cleavage of the tRNAs immediately upstream of tRNA^Leu(CUN)^ (tRNA^His^ and tRNA^Ser(AGY)^).

Our POLRMT pausing studies and insights into sequence specificity of pausing open avenues for elucidating its potential role in co-transcriptional RNA folding and processing in human mitochondria. For instance, pausing could allow for POLRMT and the nascent RNA to interact with tRNA processing enzymes (MRPP1, 2, 3, and ELAC2) and “writer” enzymes that modify the RNA^10,55^. Future *in vivo* mapping of POLRMT pauses is needed to confirm strategic transcriptional pause locations on mtDNA and their contribution to co-transcriptional regulatory events.

## Supporting information

Supporting Information - additional figures and list of oligonucleotides

## ASSOCIATED CONTENT

### Accession Codes

UniProt Accession IDs: POLRMT: O00411; TFAM: Q00059; TFB2M: Q9H5Q4, TEFM: Q96QE5; *E*.*coli* RNA Polymerase: P0A7Z4, subunit *α*; P0A8V2, subunit *β*; P0A8T7, subunit *β′*; P0A800, subunit *ω*; P00579, subunit *σ*.

## Supporting Information

The following files are available free of charge.

Supporting Information - additional figures and list of oligonucleotides (PDF)

## Author Contributions

The manuscript was written through contributions of all authors. All authors have given approval to the final version of the manuscript.

## ACKNOWLEDGMENT

We would like to thank Prof. Smita Patel (Rutgers University) and members of her lab for their generosity and protocols that aided us in this work. We would also like to thank Prof. Dmitry Temiakov (Thomas Jefferson University), Prof. Miguel Garcia-Diaz (Stony Brook University School of Medicine), Prof. Bob Landick (University of Wisconsin-Madison) for providing us with expression plasmids used in this work. We greatly appreciate Sean Reardon and Jubilee Haddasah for the purification of the various components needed for mitochondrial transcription and their valuable support throughout this work. Special thanks to Dr. Christine Stephen for her encouragement and guidance with data quantification, along with Dr. Gairika Ghosh, and Rushabh Bhakta for assisting with *E. coli* RNAP purifications. Thank you to Dr. Ravish Sharma, Dr. Mukesh Mahajan, and members of our research group for their encouragement during this study.

## ABBREVIATIONS

POLRMT: human mitochondrial RNA polymerase
mtDNA: human mitochondrial DNA
OXPHOS: oxidative phosphorylation
CSBIII, II, and I: Conserved Sequence Blocks III, II and I
TEFM: mitochondrial transcription elongation factor
EC: elongation complex
PEC: paused EC
NTP: nucleoside triphosphate
rNTP: ribonucleotide tri-phosphate

