## Supporting Information - additional figures and list of oligonucleotides for "Nucleic acid sequence determinants of transcriptional pausing by human mitochondrial RNA polymerase (POLRMT)"

An H. Hsieh, Tatiana V. Mishanina*
Department of Chemistry and Biochemistry, University of California San Diego, La Jolla, CA, USA

This pdf file includes: Supplementary Figures 1 and 2 and Table S1.

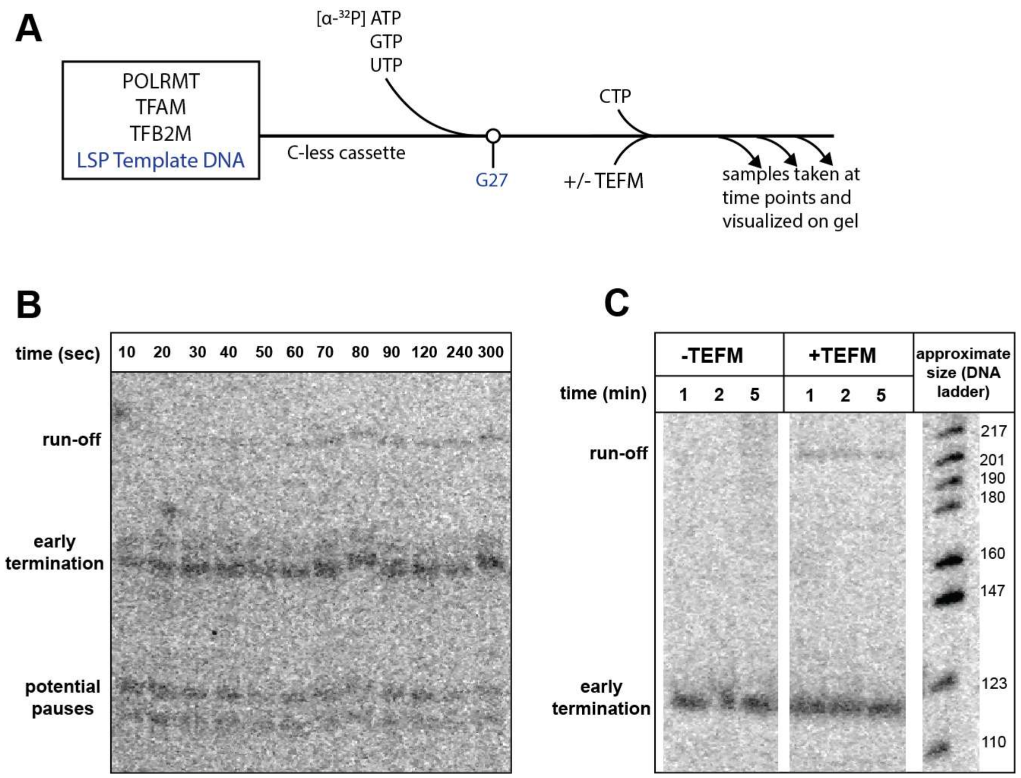


**Figure S1. Promoter-based POLRMT transcription with and without TEFM. A)** Flowchart of the promoter-based *in vitro* transcription reaction. POLRMT will begin transcription from the LSP promoter with the initiation factors TFAM and TFB2M. **B)** Time course of the promoter-based *in vitro* transcription assay, showing transcription of the full-length product over time, with early termination bands at CSBII and potential pausing sites by POLRMT. **C)** The presence of TEFM promotes processivity. Lanes of interest from reactions run side-by-side showing bands around ~120 nt that correspond to early termination at CSBII and ~220 nt for the run-off product, promoted by the addition of TEFM. RNA size was estimated against a radiolabeled DNA ladder.

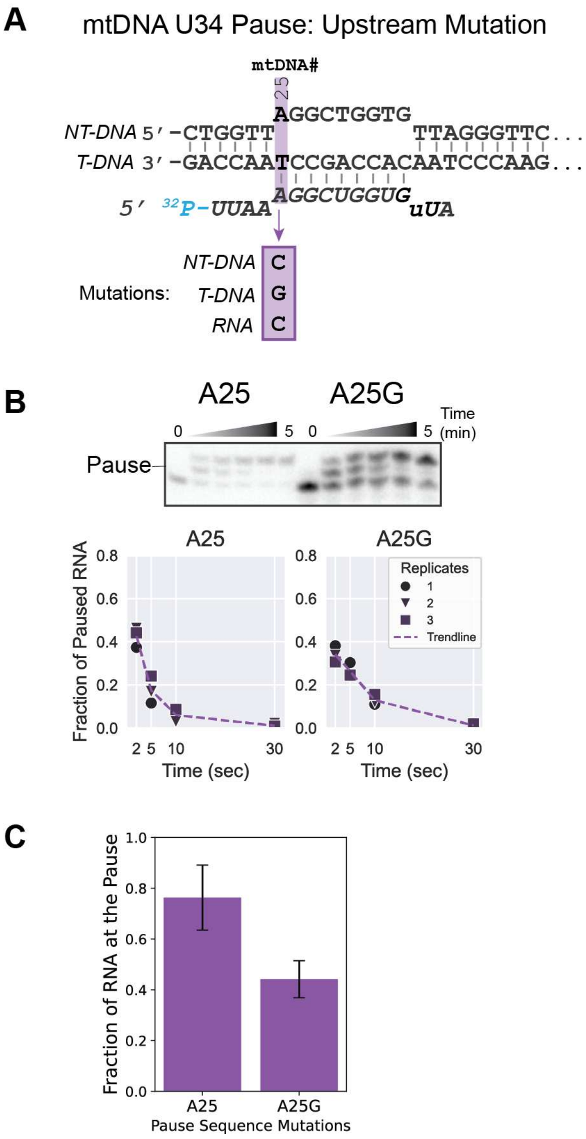


**Figure S2. Upstream mutation of the elemental POLRMT pause sequence. A)** The A25 scaffold is the same as the previous U34 scaffold except that it has a 9 nt transcription bubble to mimic the size of the *E. coli* RNAP consensus elemental scaffold. We named this new scaffold "A25" because the RNA now binds to the DNA at that position and mutated A25 to G to see the effects on pausing. **B)** We observed faint band intensity for A25 compared to A25G, but POLRMT pauses on both sequences. As seen in Fig. 1B, POLRMT prefers 3' GC-rich scaffolds, and perhaps this new scaffold starting with an A likely made it unfavorable for POLRMT binding compared to having a G at that position. **C)** The pause fraction of A25 almost decreased by half with the substitution to G, but A25G still supports POLRMT pausing.

### Table S1. Oligonucleotides Used in This Study

#### **Scaffold 1** (Fig. 1B)

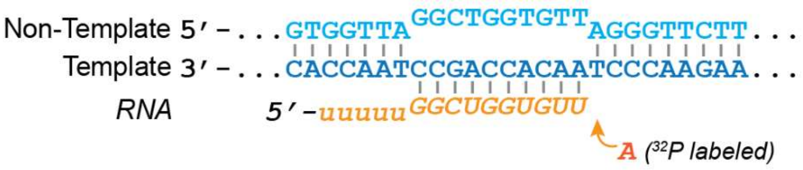


| Oligo # | Oligo name | Sequence (5'->3') | Description |
|---|---|---|---|
| 0111 | POLRMT_RNA_LSP1 | UUUUUGGCUGGUGUU | RNA that binds downstream of LSP to test for pause location in CSBIII. The 3' end is at position +35 downstream of LSP. |
| 0112 | POLRMT_TDNA_LSP1 | GCTTCTGGCCACAGCACTTAAAcACATCTCTGCCAAACCCCAAAAACAAAGAACCCTAACACCAGCCTAACCAC | Template that contains CSBIII to test for pause location. Stops before CSBII |
| 0113 | POLRMT_NTDNA_LSP1 | GTGGTTAGGCTGGTGTTAGGGTTCTTTGTTTTTGGGGTTTGGCAGAGATGTgTTTAAGTGCTGTGGCCAGAAGC | NT-DNA that is fully complementary to T-DNA that contains CSBIII |

#### **Scaffold 2 (**Fig. 1B)

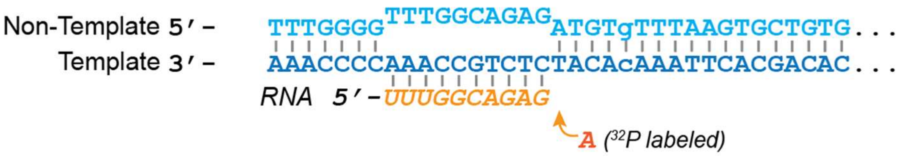


| Oligo # | Oligo name | Sequence (5'->3') | Description |
|---|---|---|---|
| 0144 | POLRMT_RNA_LSP2 | UUUGGCAGAG | RNA that binds downstream of LSP to test for pause location AFTER CSBIII. 3' end is at position +65 downstream of LSP |
| 0153 | POLRMT_NTDNA_LSP3 | TTTGGGGTTTGGCAGAGATGTgTTTAAGTGCTGTGGCCAGAAGC | To test pause after CSBIII. Stops before G-rich sequence in CSBII, to improve quality of transcripts |
| 0154 | POLRMT_TDNA_LSP3 | GCTTCTGGCCACAGCACTTAAAcACATCTCTGCCAAACCCCAAA | T-DNA that is fully complementary to 0153 |

**Scaffold 3** (SC1 RNA + Consensus elemental pause scaffold in Fig. 2)
Note: the nucleotides in green after the starting RNA are the nucleotides to be added during the experiment.

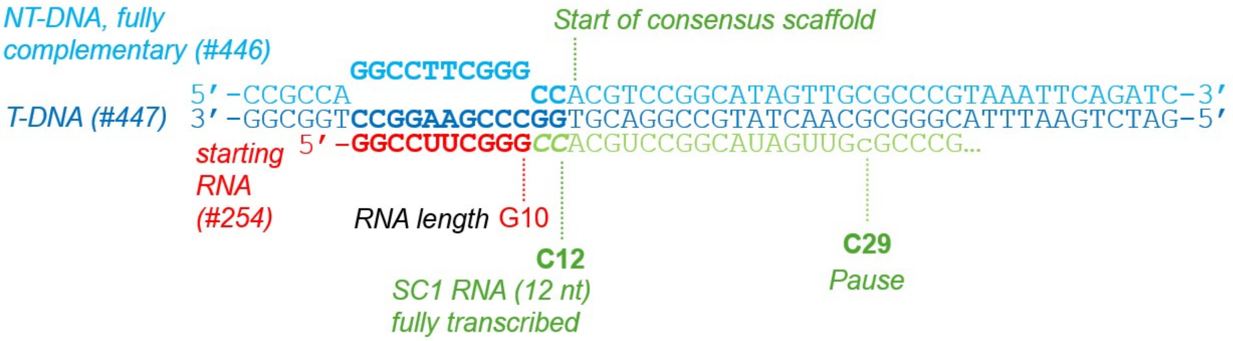


| Oligo # | Oligo name | Sequence (5'->3') | Description |
|---|---|---|---|
| 0254 | SC1_10nt | GGCCUUCGGG | RNA that forms a hairpin quickly without affecting downstream RNA structure (Merino et al. *J. Am. Chem. Soc*. 2005, CE Szyjka, EJ Strobel, *Riboregulator Design and Analysis*, 2022, CE Szyjka, EJ Strobel, *Nat Comm*, 2023). Missing the last 2 Cs to reduce $T_m$ and improve annealing to T-DNA |
| 0446 | 0446_SC1+ConsensusPause_NT-DNA | CCGCCAGGCCTTCGGGCCACGTCCGGCATAGTTGCGCCCGTAAATTCAGATC | NT-DNA for scaffold containing the SC1 RNA (#254) sequence and *E. coli* consensus elemental pause |
| 0447 | 0447_SC1+ConsensusPause_T-DNA | GATCTGAATTTACGGGCGCAACTATGCCGGACGTGGCCCGAAGGCCTGGCGG | T-DNA for scaffold containing the SC1 RNA (#254) sequence and *E. coli* consensus elemental pause |

**Scaffold 4** (SC1 RNA + downstream sequence of LSP1 in Fig. 3)

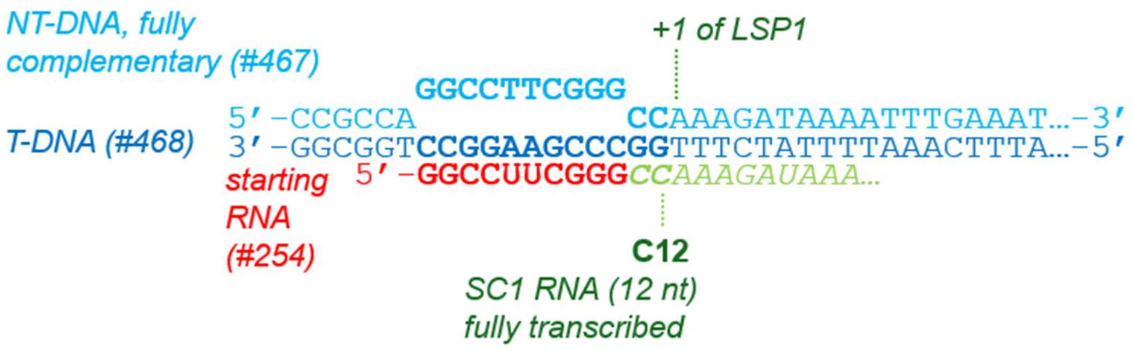


| Oligo # | Oligo name | Sequence (5'->3') | Description |
|---|---|---|---|
| 0254 | SC1_10nt | GGCCUUCGGG | RNA that forms a hairpin quickly and is used in RNA structure probing (Strobel lab). Missing the last 2 nt (CC) to reduce $T_m$ and improve annealing to T-DNA |
| 0467 | 0467_SC1_LSP_NT-DNA90nt | CCGCCAGGCCTTCGGGCCAAAGATAAAATTTGAAATCTGGTTAGGCTGGTGTTAGGGTTCTTTGTTTTTGGGGTTTGGCAGAGATGTGTT | Contains SC1 sequence, downstream LSP sequence up to +72. |
| 0468 | 0468_SC1_LSP_T-DNA90nt | AACACATCTCTGCCAAACCCCAAAAACAAAGAACCCTAACACCAGCCTAACCAGATTTCAAATTTTATCTTTGGCCCGAAGGCCTGGCGG | T-DNA that is fully complementary to the NT-DNA. |

**Scaffold 5** (U34 scaffold Figs. 5 and 6)

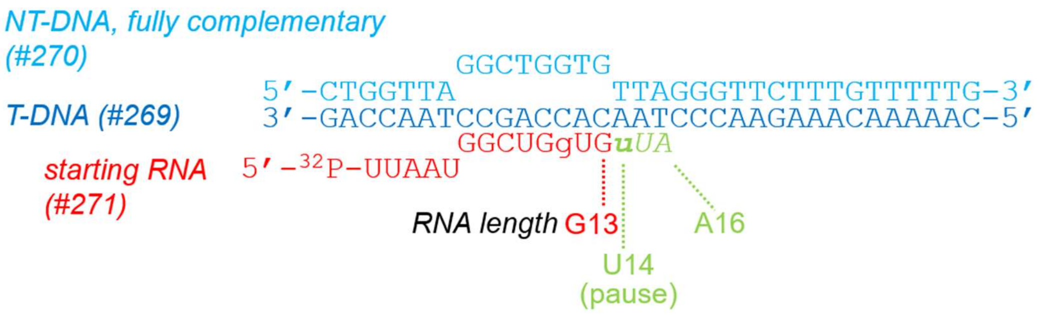


| Oligo # | Oligo name | Sequence (5'->3') | Description |
|---|---|---|---|
| 0269 | T-DNA_LSP _pause34 | CAAAAACAAAGAACCC TAACACCAGCCTAACC AG | T-DNA containing sequence downstream of LSP for pause at +34 |
| 0270 | NT-DNA_LSP _pause34 | CTGGTTAGGCTGGTGT TAGGGTTCTTTGTTTTT G | NT-DNA containing sequence downstream of LSP for pause at +34. Fully complementary to the TDNA. |
| 0271 | 1_nt_before_ pause34_RNA | UUAAUGGCUGgUG | RNA that ends 1 nt before the pause at +34 downstream of LSP. 8 nt will bind to the T-DNA (13 nt total) The first 5 nucleotides were randomly chosen and do not bind to the T-DNA. |

**Scaffold 7** (U34G scaffold in Fig. 6)

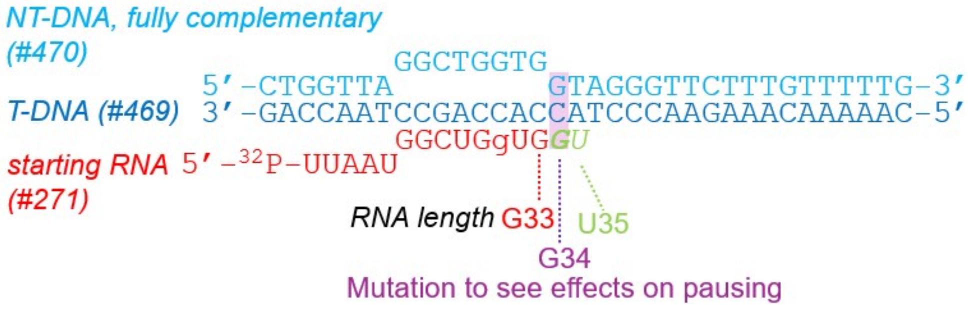


The RNA is the same in the U34 Scaffold above (#0271)

| Oligo # | Oligo name | Sequence (5'->3') | Description |
|---|---|---|---|
| 0469 | 0469_T-DNA_LSP_ pauseU34G | CAAAAACAAAGAAC CCTACCACCAGCCT AACCAG | T-DNA same as #0269 in the U34 Scaffold except has mutation from U34>G to test for effect on the pause. |
| 0470 | 0470_NT-DNA_LSP_ pauseU34G | CTGGTTAGGCTGGT GGTAGGGTTCTTTG TTTTTG | NT-DNA same as #0270 in the U34 Scaffold except has mutation from U34>G to test for effect on the pause. |

**Scaffold 8** (U34C scaffold in Fig. 6)

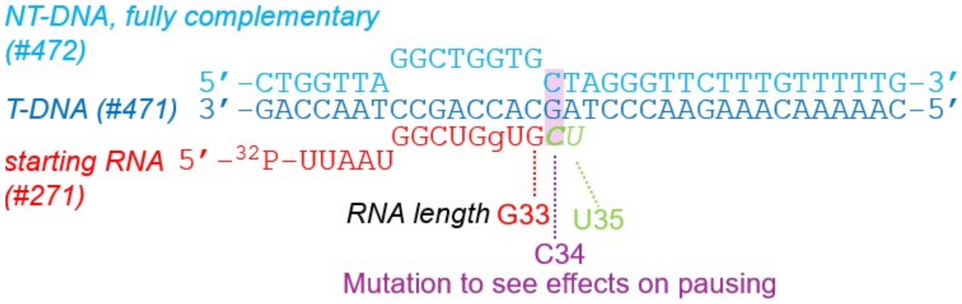


The RNA is the same in the U34 scaffold above (#0271)

| Oligo # | Oligo name | Sequence (5'->3') | Description |
|---|---|---|---|
| 0471 | 0471_T-DNA_LSP_pauseU34C | CAAAAACAAAGAACCCTAGCACCAGCCTAACCAG | T-DNA same as #0269 except has mutation from U34>C to test for effect on the pause. |
| 0472 | 0472_NT-DNA_LSP_pauseU34C | CTGGTTAGGCTGGTGCTAGGGTTCTTTGTTTTTG | NT-DNA same as #0270 except has mutation from U34>C to test for effect on the pause. |

**Scaffold 9** (U35G scaffold in Fig. 6)

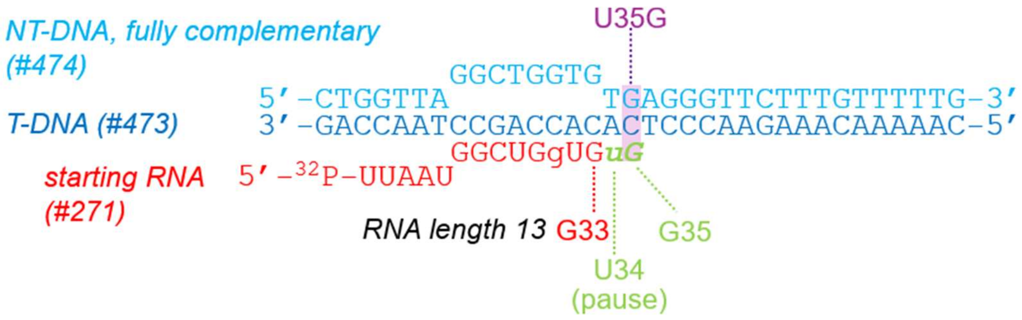


The RNA is the same in the U34 scaffold above (#0271)

| Oligo # | Oligo name | Sequence (5'->3') | Description |
|---|---|---|---|
| 0473 | 0473_T-DNA_LSP_incomingU35G | CAAAAACAAAGAACCCTCACACCAGCCTAACCAG | T-DNA same as #0269 except has mutation from the incoming U35>G to test for effect on the pause. |
| 0474 | 0474_NT-DNA_LSP_incomingU35G | CTGGTTAGGCTGGTGTGAGGGTTCTTTGTTTTTG | NT-DNA same as #0270 except has mutation from the incoming U35>G to test for effect on the pause. |

**Scaffold 10** (U35C scaffold in Fig. 6)

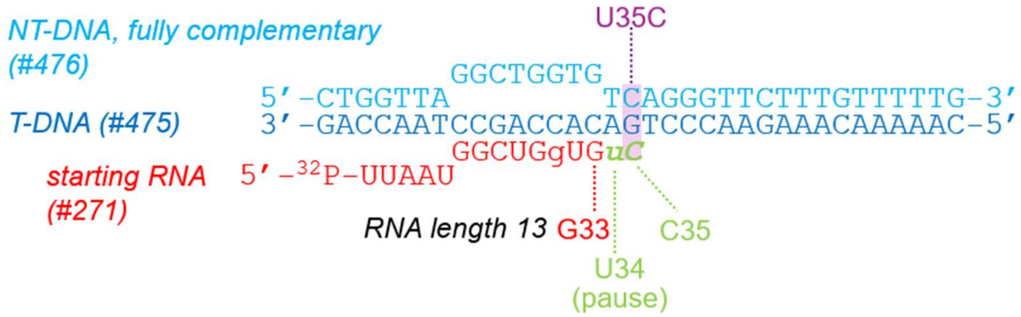


The RNA is the same in the U34 scaffold above (#0271)

| Oligo # | Oligo name | Sequence (5'->3') | Description |
|---|---|---|---|
| 0475 | 0475_T-DNA_LSP_incomingU35C | CAAAAACAAAGAACCCTGACACCAGCCTAACCAG | T-DNA same as #0269 except has mutation from the incoming U35>C to test for effect on the pause |
| 0476 | 0476_NT-DNA_LSP_incomingU35C | CTGGTTAGGCTGGTGTCAGGGTTCTTTGTTTTTG | NT-DNA same as #0270 except has mutation from incoming U35>C to test for effect on the pause |

**Scaffold 11**(G33U scaffold in Fig. 6)

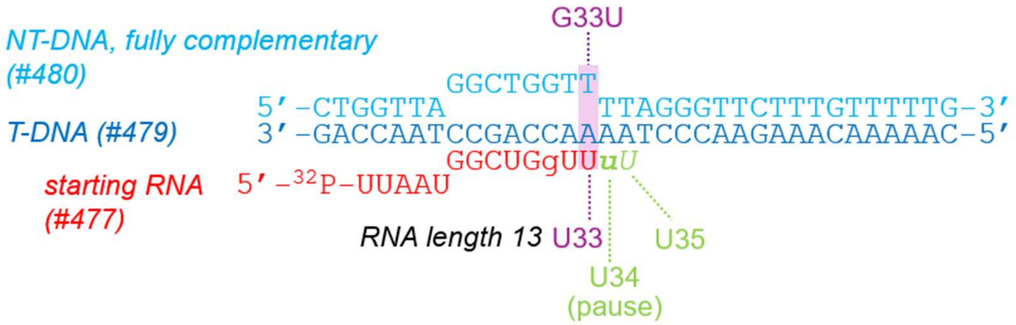


| Oligo # | Oligo name | Sequence (5'->3') | Description |
|---|---|---|---|
| 0477 | 0477_1ntbeforePause34G33U | UUAAUGGCUGgUU | Mutated version of #271 from G33 to U. RNA that ends 1 nt before the pause at 34 nt downstream of LSP. The first 5 nt are random and the next 8 nt will bind to the T-DNA. RNA length is 13 nt |
| 0479 | 0479_T-DNA_LSP_pauseG33U | CAAAAACAAAGAACCCTAAAACCAGCCTAACCAG | T-DNA same as #0269 except has mutation G33>U to test the role of the -2 position on the pause. |

| | | | |
|---|---|---|---|
| 0480 | 0480_NT-DNA_LSP_pauseG33U | CTGGTTAGGCTGGTTTTAGGGTTCTTTGTTTTTG | NT-DNA same as #0270 except has mutation G33>U to test the role of the -2 position on the pause. |

**Scaffold 12** (G33C scaffold in Fig. 6)

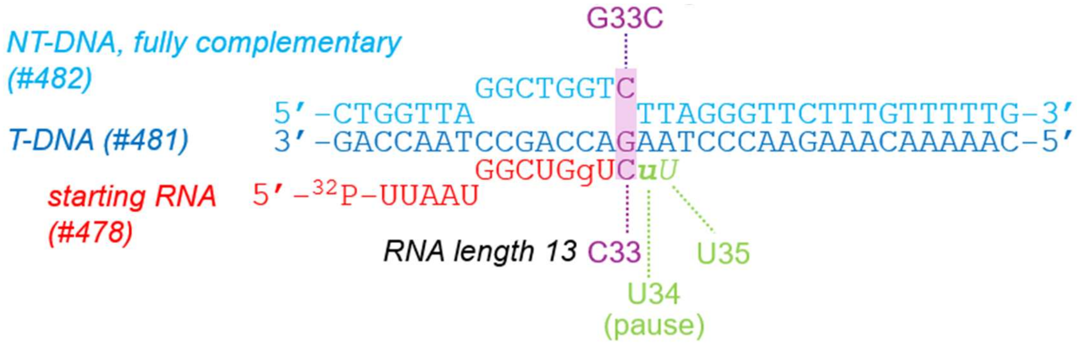


| Oligo # | Oligo name | Sequence (5'->3') | Description |
|---|---|---|---|
| 0478 | 0478_1ntbeforePause34G33C | UUAAUGGCUGgUC | Mutated version of #271 from G33 to C. RNA that ends 1 nt before the pause at 34 nt downstream of LSP. The first 5 nt are random and the next 8 nt will bind to the T-DNA. RNA length is 13 nt. |
| 0481 | 0481_T-DNA_LSP_pauseG33C | CAAAAACAAAGAACCCTAAGACCAGCCTAACCAG | T-DNA same as #0269 except has mutation G33>C to test the role of the -2 position on the pause |
| 0482 | 0482_NT-DNA_LSP_pauseG33C | CTGGTTAGGCTGGTCTTAGGGTTCTTTGTTTTTG | NT-DNA same as #0270 except has mutation G33>C to test the role of the -2 position on the pause |

**Scaffold 13** (A25 scaffold in Fig. S2)

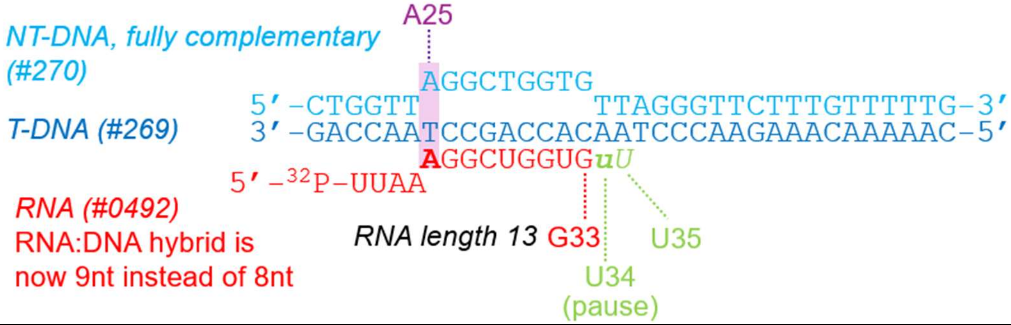


| Oligo # | Oligo name | Sequence (5'->3') | Description |
|---|---|---|---|
| 0492 | 0492_RNA_LSP_-10WT_A25 | UUAAAGGCUGGUG | RNA to make scaffold with T-DNA 0269 and NT-DNA 0270 which now forms a 9nt RNA:DNA hybrid |

| | | | |
|---|---|---|---|
| 0269 | T-DNA_LSP_pause34 | CAAAAACAAAGAACCCTAACACCAGCCTAACCAG | Same T-DNA used in the U34 scaffold above (Scaffold 5). |
| 0270 | NT-DNA_LSP_pause34 | CTGGTTAGGCTGGTGTTAGGGTTCTTTGTTTTTG | Same NT-DNA used in the U34 scaffold above (Scaffold 5). |

**Scaffold 14** (A25G scaffold as seen in Fig. S2)

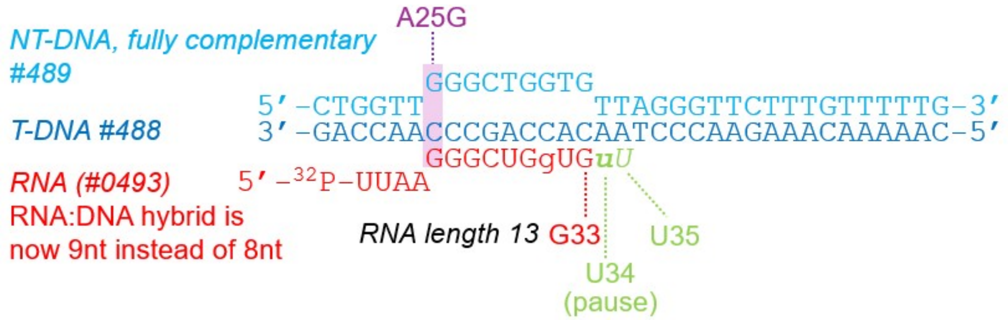


| Oligo # | Oligo name | Sequence (5'->3') | Description |
|---|---|---|---|
| 0493 | 0493_RNA_LSP_-10mutation_A25G | UUAAGGGCUGGUG | RNA to make scaffold with T-DNA 0488 and NT-DNA 0489 which now forms a 9nt RNA:DNA hybrid with mutation at -10 (A25G) |
| 0488 | 0488_T-DNA_LSP_-10mutation_A25G | CAAAAACAAAGAACCCTAACACCAGCCCAACCAG | T-DNA same as #0269 except has mutation A25>G to test the role of the -10 position on the pause |
| 0489 | 0489_NT-DNA_LSP_-10mutation_A25G | CTGGTTGGGCTGGTGTTAGGGTTCTTTGTTTTTG | NT-DNA same as #0270 except has mutation A25>G to test the role of the -10 position on the pause |
